# cfDecon: Accurate and Interpretable methylation-based cell type deconvolution for cell-free DNA

**DOI:** 10.1101/2025.02.11.637663

**Authors:** Yixuan Wang, Jiayi Li, Jingqi Li, Shen Yang, Yuhan Huang, Xinyuan Liu, Yimin Fan, Irwin King, Yumei Li, Yu Li

## Abstract

Cell-free DNA (cfDNA) analysis has emerged as a powerful tool for noninvasive diagnostics and monitoring of aberrant methylation. However, computational deconvolution of cfDNA remains challenging due to the complexity of methylation data, the diverse cellular compositions in cfDNA mixtures, and the limited interpretability of current methods. We present cfDecon, an efficient deep-learning framework for cfDNA deconvolution at the resolution of individual reads. cfDecon employs a multichannel autoencoder core module and an iterative refinement process to estimate cell-type proportions and generate condition-aware cell-type-specific methylation profiles. We rigorously evaluate cfDecon through comprehensive simulation experiments, including scenarios with normal cellular compositions, rare cell types, and unknown cell types, where cfDecon consistently outperforms previous state-of-the-art methods in deconvolution. Using strict data separation and careful control for potential data leakage, cfDecon shows scalability and applicability to large-scale studies by raising Lin’s Concordance Correlation Coefficient (CCC) over 33% on an atlas-level reference dataset. We further validate cfDecon’s utility in disease diagnosis on two clinical datasets, Amyotrophic Lateral Sclerosis (ALS) and Hepatocellular Carcinoma (HCC). cfDecon raises the performance of disease detection from 0.53 to 0.79 compared to existing methods on the ALS dataset and 0.55 to 0.77 on the HCC dataset. The capacity of cfDecon to predict condition-specific DNA methylation patterns across cell types offers a valuable framework for studying differentially methylated CpGs. Furthermore, integrating cfDecon with enrichment analysis facilitates the exploration of functional variations in CpGs across different conditions. cfDecon represents a significant advancement in cfDNA analysis, offering improved accuracy, interpretability, and potential for personalized medicine applications. The codebase is available at https://github.com/Susanxuan/cfDecon.

## 1 Introduction

Short DNA fragments released into the blood plasma are known as cell-free DNA (cfDNA) [27,64]. It primarily originates from cellular apoptosis throughout the body and has emerged as a powerful tool for monitoring aberrant methylation and noninvasive diagnostics [25,9]. Delineating the role of methylation alterations in disease pathophysiology necessitates precise mapping of epigenetic landscapes in both healthy and diseased cells; however, this endeavor is complicated by heterogeneous cell type compositions and intra-cell-type variability in primary tissues [60,48]. Cell-of-origin deconvolution of cfDNA with DNA methylation patterns, facilitating applications in prenatal testing [20], early detection of tissue damage across multiple organs simultaneously [19], cancer and autoimmune disorder diagnosis [71,42], and monitoring of treatment response and disease recurrence [23,37]. The capacity for noninvasive, whole-body wellness profiling renders cfDNA analysis a promising frontier in personalized medicine.

However, computational deconvolution of cfDNA remains a challenge. The first issue arises from the various types of methylation data ranging from affinity-enrichment methods capturing overall regional methylation levels to bisulfite-conversion, enzymatic conversion, and third-generation long-read sequencing technologies providing single-CpG resolution [1,31,81]. Conventional cfDNA deconvolution approaches typically utilize aggregated methylation levels at each CpG site, potentially overlooking crucial information by assuming independence between individual CpG sites despite considering regional correlations [7,29]. Secondly, though cell-type proportion prediction in cfDNA datasets has been extensively studied, methods capable of generating cell-type-specific DNA methylation profiles remain scarce [83,55]. This limitation restricts the interpretability of current decomposition approaches and hinders the detection of methylation aberrations between healthy and disease states. Thirdly, the complexity of cfDNA mixtures, often comprising more than 35 cell types in reference atlases [49], necessitates the development of deep learning-based tools. Traditional methods struggle to capture the heightened nonlinear properties of such diverse cellular compositions, underscoring the need for more sophisticated computational approaches.

Several computational methods, including both statistical and deep learning methods, have been developed to dissect cfDNA datasets. These approaches can be broadly categorized into two classes based on their utilization of methylation data: CpG site-based and read-based methods. The former category, which includes techniques such as CelFiE [7], UXM [49], CelFiE-ISH [70], and cfSort [44], employs the mean methylation level across all sequencing reads per CpG site as model input. While these site-based approaches have demonstrated efficacy in elucidating the tissue/cell origin of cfDNA, they potentially sacrifice sensitivity and specificity by disregarding the correlation between adjacent CpG sites. This hypothesis has been corroborated in the context of tumor fraction estimation, a problem with similar settings [45,43,52]. In contrast, the only read-based method now available is CelFEER [33]. This method employs an expectation-maximization (EM) algorithm to estimate parameters within a Bayesian model of cfDNA mixture cell-type proportions, building upon the foundation laid by CelFiE. Fig. 1 **a** illustrates the input formats for both CpG site-based and read-based methods, with read-based processing providing higher resolution through five-channel encoding. A tumor-derived read can be distinguished from other reads more easily by comparing read-based encoding instead of CpG site averages as shown in Fig. 1 **b**. Notably, a significant limitation across all existing methodologies is their failure to generate condition-aware cell-type-specific methylation profiles.

**Fig. 1:**
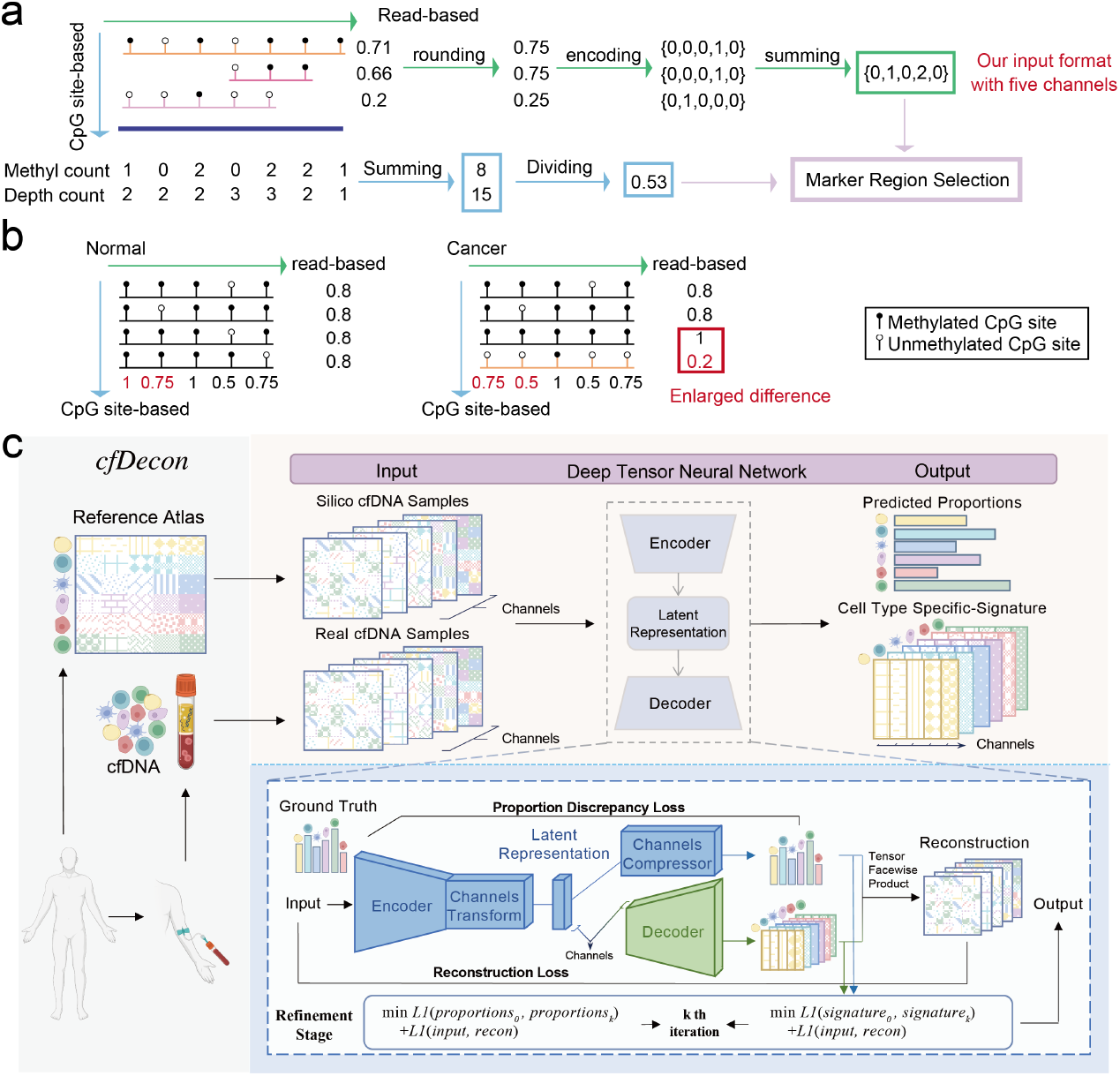
Overview of cfDecon. **a**. Formatting of the input for CpG site-based and read-based methods. Three partially methylated reads aligning to a 500 bp region are depicted. For CpG site-based method, the input is the mean methylation level across all sequencing reads per CpG site. For read-based methods, the read averages are rounded to discrete values 0, 0.25, 0.5, 0.75, and 1, encoded in a one-hot format, and summed to represent counts for each methylation channel within a 500 bp region. **b**. Example showing improved tumor read detection using read-based versus CpG site-based analysis [33]. **c**. Detailed overview of the cfDecon, illustrating the data collection process on the left, which includes both silico data and real cfDNA samples, and the right side depicts the core components of cfDecon within a multichannel autoencoder framework, showing how input data is processed to yield estimated cell-type proportions and cell-type-specific methylation signature.

To overcome these limitations, we propose an accurate, efficient, and interpretable deep-learning algorithm, cfDecon, for read-based cfDNA deconvolution. In particular, cfDecon adopts a low-rank structure to avoid the heavy redundancy of multichannel autoencoder. cfDecon takes cfDNA methylation data as input and estimates cell-type proportions and cell-type-specific signature profiles. To generate condition-aware cell-type-specific signatures, the model iteratively adapts its parameters to incoming cfDNA data through a refinement stage, alternating between decoder optimization for signature generation and encoder adjustment for proportion prediction. These architectural innovations ensure superior performance rather than relying on overfitting. On top of that, cfDecon provides an integrated pipeline for cfDNA component analysis and methylation aberration detection, enabling comprehensive cfDNA-based disease diagnostics.

Through rigorous data partitioning and systematic prevention of data leakage, we evaluate cfDecon through extensive simulations across typical cellular compositions, rare cell populations, and missing cell type scenarios. Our results consistently demonstrate cfDecon’s superior performance over existing state-of-the-art deconvolution methods. cfDecon’s scalability and applicability to large-scale studies are particularly noteworthy, demonstrating over 33% improvement in Lin’s Concordance Correlation Coefficient (CCC) on an atlas-level reference dataset with 35 distinct cell types. Clinical validation using disease-specific datasets shows enhanced disease detection capabilities, improving diagnosis accuracy from 0.53 to 0.79 for Amyotrophic Lateral Sclerosis (ALS) and from 0.55 to 0.77 for Hepatocellular Carcinoma (HCC). Beyond accurate deconvolution, cfDecon provides a framework for analyzing condition-specific DNA methylation patterns and enables functional CpG variation analysis through enrichment analysis. We demonstrate that cell-typespecific signatures generated by cfDecon can differentiate between disease and healthy samples with biological significance. These capabilities establish cfDecon as a significant advancement in cfDNA analysis, offering superior accuracy and enhanced interpretability for potential personalized medicine applications.

## 2 Methods

### 2.1 Problem Definition and Target Data Format

To elucidate our model, it is crucial to define the problem in advance. We aim to deconvolute cell-free DNA (cfDNA) methylation data to determine the relative contributions of different cell types. Building on prior studies [7,33], we assume that the methylation profile of cfDNA represents a linear combination of methylation profiles from distinct cell types. Given *t* cell types and *n* samples, our goal is to estimate the cell-type proportions contributing to the cfDNA methylation profile and predict the cell-type-specific signature to provide more insights. Mathematically, this can be represented as:

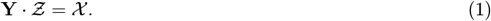

Here χ ∈ ℝ ^*n×m×c*^ is the target cfDNA data, Ƶ ∈ ℝ ^*t×m×c*^ is the cell-typespecific methylation signature tensor (*m* represents the CpG marker regions and *c* denotes the channel dimension), and **Y ∈** ℝ ^*n×t*^ is the cell-type proportions matrix.

Studies demonstrate that methylation averages across entire reads provide better detection of tumor-derived DNA fragments compared to individual CpG sites [7,29,33] (Fig. 1 **b**). Following CelFEER’s approach [33], we replace individual CpG site counts with five counts per 500 bp region. For each DNA fragment with at least three CpG sites, we calculate methylation density as the ratio of methylated to total CpG sites, discretizing it to the nearest quartile value 0, 0.25, 0.5, 0.75, and 1. This process generates a one-hot encoded vector with five frequency values (c=5) per region (Fig. 1 **a**). Considering computational constraints, we select informative regions by determining cell-type pair distances based on read-level methylation averages following [33] corresponding to the discriminative markers for the model input.

### 2.2 Multichannel Autoencoder Framework

We develop a deep tensor neural network within a multichannel autoencoder framework for cfDNA deconvolution named cfDecon (Fig. 1 **c**). cfDecon integrates cell-type-specific DNA methylation profiles across *t* cell types and *m* CpG marker regions, along with *n* simulated or real cfDNA samples *X* . To enhance interchannel interactions, we implement a channel transform function *ϕ*(·) to get the latent representation. Then the predicted cell-type proportions matrix Ŷ is obtained through the channel compressor *ψ*(·). Sequentially, the decoder is expected to reconstruct the cfDNA tensor 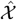 with the predicted proportions matrix Ŷ. This makes the decoder function like the cell-type-specific signature tensor 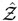, thus 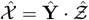. To achieve interpretability, our decoder is designed without activation layers or biases inspired by our previous work [11], which is only the regularized value of the dot product of weight matrices. This process can be formulated as:

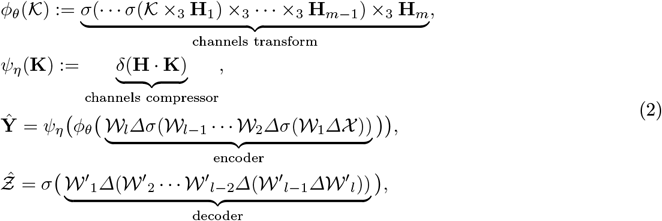

where Δ is the tensor face-wise product defined by *𝒞* = *𝒜*Δ *ℬ* ⇔ *𝒞*^(*k*)^ = *𝒜* ^(*k*)^ *ℬ*^(*k*)^ [32], 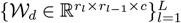 are some factor tensors of encoder, *l* is the number of layer, *σ*(·) denotes the LeakyReLU activation function, and 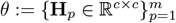 are learnable parameters of the channels transform. Additionally, · denotes matrix product, *×*3 denotes the mode-3 tensor-matrix product [34], 𝒦 and **K** are introduced as an illustrative variable to demonstrate the function’s behavior, *η* := {**H** ∈ R^*tc×c*^} are learnable parameters of the channel’s compressor, δ(·) denotes the Softmax function, and 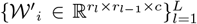 are factor tensors of decoder. Considering the multichannel autoencoder may suffer from significant redundancy, we also adopt a low-rank structure to enhance the efficiency of our model (see quantitative low-rank analysis in Supplementary S1.1).

### 2.3 Training Method

We generate in silico data with pre-defined proportions **Y** for model training and validation (Supplementary S1.4). Our objectives include both cell-type proportion prediction and the reconstruction of cfDNA. The encoder predicts cell-type proportions **Ŷ** by minimizing the mean absolute error (MAE) with ground truth proportions. The decoder optimizes cell-type-specific methylation signatures 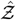 to reconstruct cfDNA 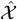 through matrix multiplication with **Ŷ** . The complete loss function is:

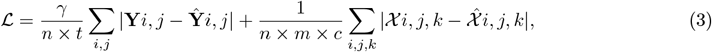

where *γ* denotes the hyperparameter. cfDecon employs a two-stage training protocol with initial training and refinement, where the refinement stage adapts parameters to novel test data through iterative optimization:

#### Optimize the decoder

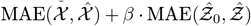 ·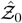, where Ŷ_0_ represents the signature tensor after initial training, or true signature tensor Ƶin real data;

#### Optimize the encoder

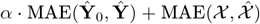, where Ŷ_0_ represents the proportions matrix after initial training, and *α, β* are hyperparameters.

The rationale underlying this iterative approach is to enhance model generalizability through data-specific adaptation. The standardized training procedure can be found in Supplementary S1.2. While convergence is not guaranteed, we constrain the adjusted parameters to remain proximate to their initial values. The decoupled training of encoder and decoder components promotes convergence stability. Multiple iterations of this optimization process typically yield parameters that effectively accommodate novel data distributions. More detailed architecture and hyperparameter settings can be found in Supplementary S1.3.

## 3 Results

### 3.1 Extensive Assessment Using Multi-Faceted in Silico cfDNA Data

Current technological limitations preclude accurate determination of cellular origins in cfDNA samples, which fundamentally impedes effective model training and evaluation due to scarce ground-truth cell-type composition data. To mitigate this limitation, we generate in silico cfDNA data from cell-type DNA methylation maps [14,49] with pre-defined cell-type proportions to benchmark existing deconvolution tools. We explain the detailed in silico cfDNA Data generation process and evaluation metrics in Supplementary S1.4 and S1.6. Given the absence of established methods, we design a deep learning-based approach, Baseline-P, as a benchmark in Supplementary S1.7.

Using the whole-genome shotgun bisulfite sequencing (WGBS) reference from ENCODE [14], we define three deconvolution scenarios. The “normal” scenario employs randomly generates cell-type proportions with sparse distributions. The “rare” scenario simulates low-abundance cell populations by assigning certain cell types fractions below 3%. The “missing” scenario generates pseudo-cfDNA data using the full reference while intentionally omitting some cell types during proportion inference. In the “normal” scenario in Fig. 2 **a**, cfDecon demonstrates superior performance across different sparsity levels (0.1, 0.3, 0.5), consistently achieving the best L1 scores, which are 0.0031, 0.0044, and 0.0069, respectively, and the highest CCC scores, which are 0.9978, 0.9968, and 0.9954, with smaller interquartile ranges. Baseline-P shows intermediate performance, while CelFEER exhibits the worst L1 scores, which are 0.0539, 0.0538, and 0.0624, and the lowest CCC scores of 0.4153, 0.4972, and 0.5503. Our model outperforms CelFEER with a p-value less than 0.05 based on the two-tailed paired t-test. In the “rare” scenario in Fig. 2 **b**, cfDecon maintains its superior performance with the worst L1 scores of 0.0047, 0.0053, and 0.0064, and the highest CCC scores of 0.9964, 0.9961, and 0.9959. The performance gap between cfDecon and other methods appears more pronounced in this challenging scenario, particularly for L1 scores where our model significantly outperforms others. In the “missing” scenario in Fig. 2 **c**, all methods show declining performance as the number of missing cell types increases from 0 to 2. However, cfDecon and our constructed Baseline-P maintain higher Pearson correlation coefficients compared to CelFEER, especially when two cell types are missing from the reference. Specifically, the cell-type proportion-based correlation score is 0.9579 for our model, 0.9487 for Baseline-P, and 0.7969 for CelFEER. To provide a more comprehensive evaluation, we conduct additional analyses as shown in Supplementary Fig. S9, which tracks the performance degradation patterns of cfDecon as cell types are progressively removed from the reference. In Supplementary Fig. S10, we examine the relationship between factorized residuals and true proportions for cell types in the pseudobulbs. The NMF factors show strong correlations with the proportions of missing cell types (1-3 missing cell types), demonstrated by high Pearson correlation coefficients and low RMSE values. This analysis validates our method’s capability to accurately capture missing cell type information through the learned latent factors.

**Fig. 2:**
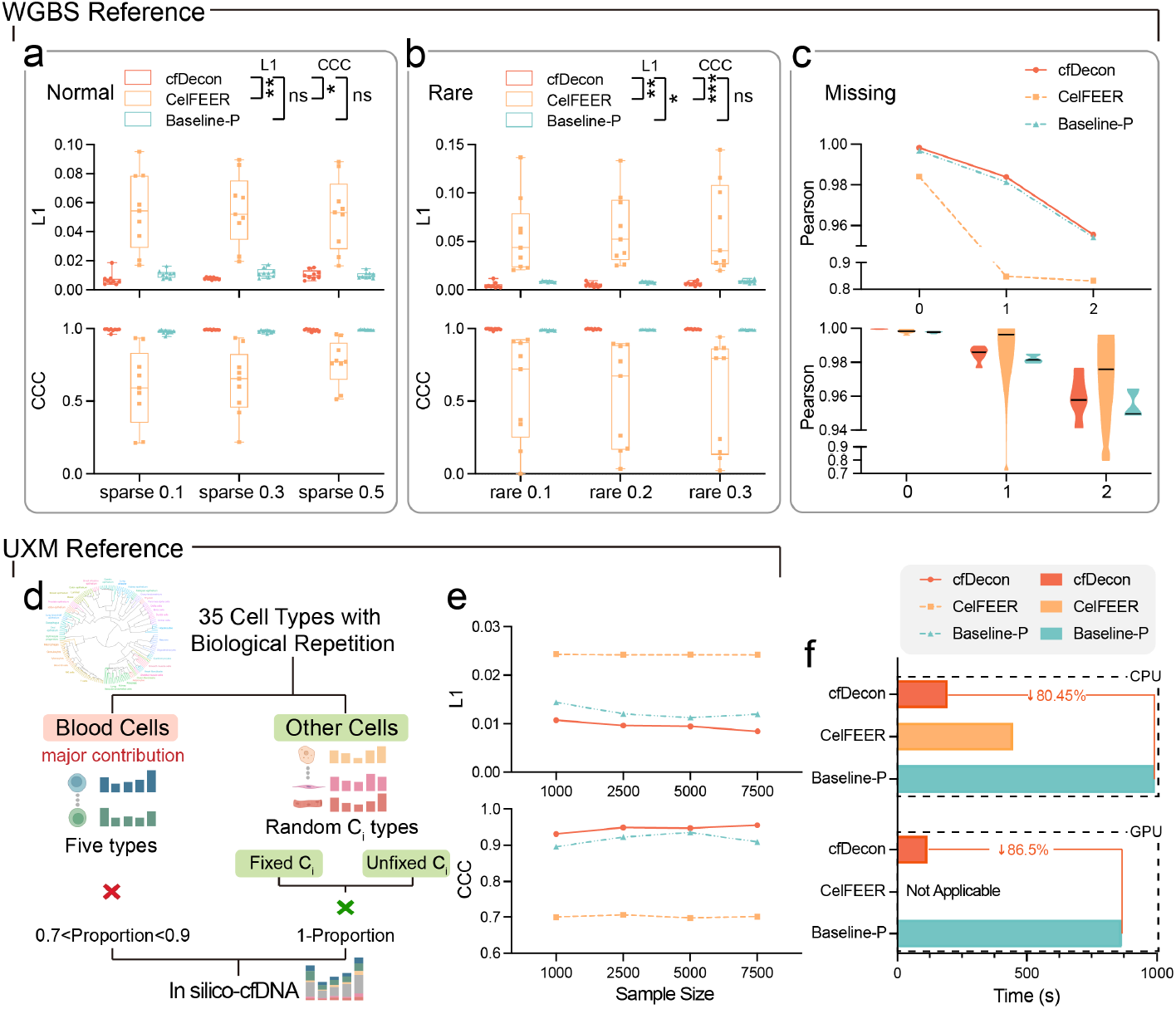
Comprehensive evaluation of deconvolution methods on in silico cfDNA data. ‘Baseline-P’ stands for our constructed deep learning-based baseline for proportion prediction. **a**,**b**. Performance comparison in normal and rare cell type scenarios with varying sparsity and rarity levels using WGBS reference data. **c**. Assessment of Pearson’s correlation performance with missing cell types. **d**. Illustration of the proportion setup for in silico cfDNA generation using the UXM reference dataset with 35 cell types. From a reference set of 35 cell types with biological replicates, this workflow simulates cfDNA mixtures by combining dominant blood cell contributions with smaller proportions from randomly selected non-blood cells, using either a fixed mode (same cell type numbers across all samples) or an unfixed mode (randomly selected cell type numbers per sample). **e**. Performance comparison across different sample sizes using the UXM reference dataset in the unfixed mode. **f**. Computational time comparison on 1000 samples for CPU and GPU implementations.

The recently published high-quality WGBS cell-type atlas [49] enables us to evaluate our method’s scalability across diverse cell types and tissues. To generate biologically meaningful in silico cfDNA data, we implement a simulation strategy that mirrors real-world cfDNA compositions (Fig. 2 **d**). We allocate 70-90% of the total composition to five major blood cell types, with the remaining proportion distributed among at least nine randomly selected cell types from 30 non-blood categories for each sample. Our simulations support both fixed mode (randomly select nine cell types across all samples) and unfixed mode (select random numbers of cell type per sample). Using this simulation framework, we analyze the effect of sample size on deconvolution performance in Fig. 2 **e**. Baseline-P peaks at 5000 samples before showing a slight decline at 7500 samples, and CelFEER shows the least favorable results. cfDecon consistently outperforms both CelFEER and Baseline-P across all sample sizes and maintains superior performance with slight improvements at larger sample sizes, demonstrating its effectiveness for large-scale deconvolution tasks.

Computational efficiency represents a critical factor in real-world cfDNA analysis, particularly for achieving scalability in large-scale applications. We evaluate the computational performance using 1000 samples for WGBS reference data. Benchmarking results demonstrate that cfDecon achieves an 80.45% reduction in CPU processing time and an 86.57% reduction in GPU processing time compared to the baseline-P method. While CelFEER exhibits intermediate CPU performance, it lacks GPU compatibility. The GPU acceleration capability of cfDecon significantly enhances its throughput potential, making it particularly suitable for large-scale cfDNA deconvolution tasks that require rapid processing of substantial data volumes.

### 3.2 Efficient Prediction of the Cell-type-specific Signatures at High-resolution

cfDecon advances beyond existing methods by generating context-aware cell-type-specific DNA methylation profiles alongside cell fraction predictions. This enables cfDecon to train on healthy data while maintaining accurate predictions under pathological conditions. cfDecon operates in two modes (Fig. 3 **a**): an “overall” mode that generates a single signature matrix for the entire dataset, and a “high-resolution” mode that produces sample-specific cell-type signatures (Supplementary S1.5). To address the computational challenges of multi-channel signature prediction, cfDecon employs tensor processing and low-dimensional projection, significantly reducing computational complexity while preserving low-rank properties (Supplementary S1.1). For a better benchmark, we develop a baseline signature generation method, Baseline-S, with implementation details demonstrated in Supplementary S1.7.

**Fig. 3:**
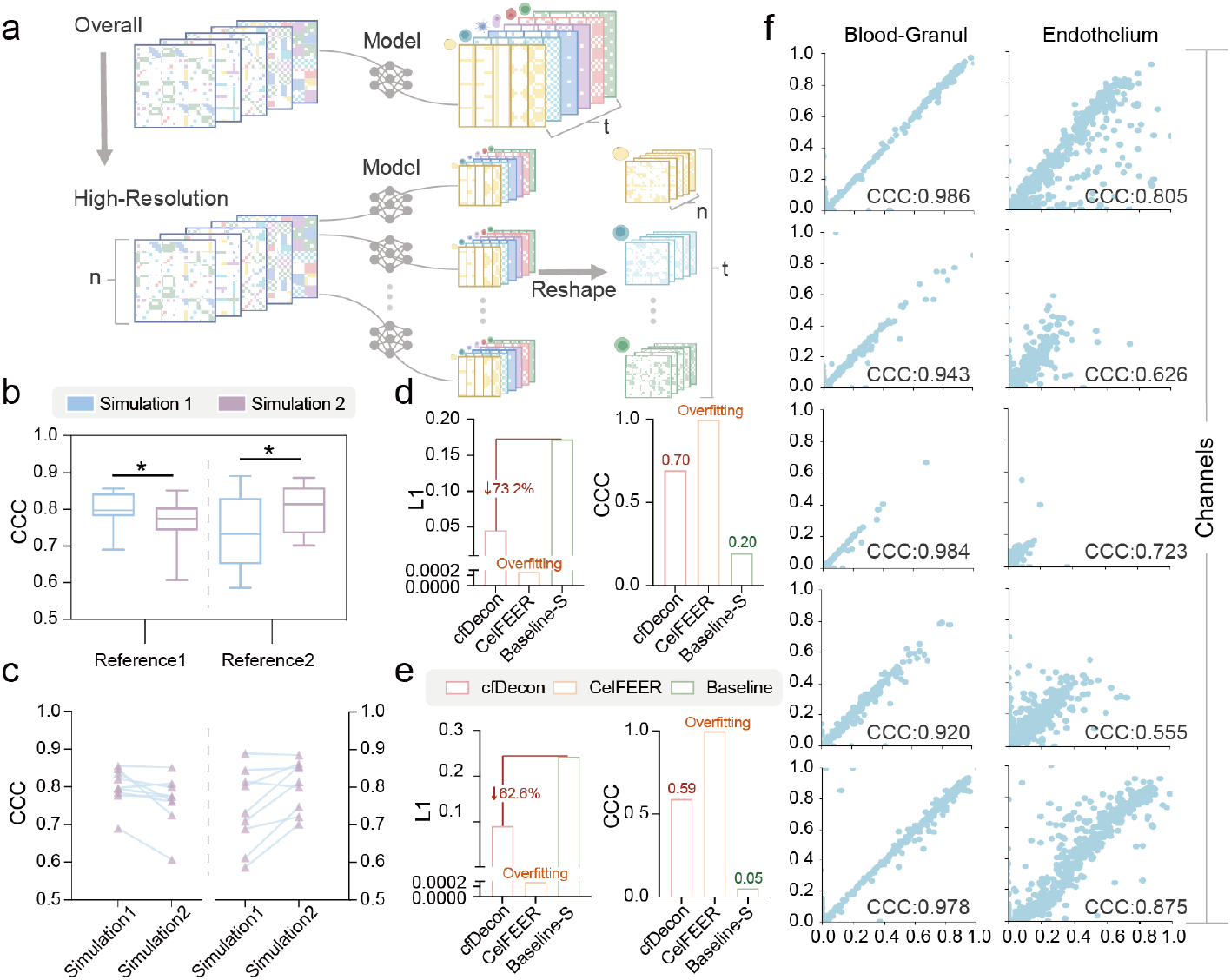
Generation of cell-type-specific signature profiles. **a**. Schematic overview of overall and high-resolution modes for predicting cell-type-specific methylation signature. **b**. Analysis of the context-aware signatures in high-resolution mode, where “Simulation 1” and “Simulation 2” correspond with the in silico data generated from “Reference 1” and “Reference 2” respectively. **c**. The CCC values for individual cell type signatures in the context-aware analysis. **d**,**e**. Signature performance comparison of cfDecon, CelFEER, and Baseline-S on the fixed **d**. and unfixed versions **e**. of the UXM reference datasets with 5000 samples in the overall mode. **f**. Detailed visualization for individual channels in the fixed version of the UXM reference dataset under the 5000 training samples, with corresponding CCC values.

To evaluate cfDecon’s context-aware signature generation capability, we design a high-resolution mode experiment using two WGBS reference datasets with biological replicates. We test signatures derived from deconvolving “Simulation 1” and “Simulation 2” using both “Reference 1” and “Reference 2”. The results demonstrate dual adaptability: signatures from “Simulation 1” show higher similarity to “Reference 1” than those from “Simulation 2”, and vice versa (Fig. 3 **b**). This pattern achieves statistical significance under two-tailed t-test analysis. Additionally, cell-type signatures consistently demonstrate higher CCC values when matched with their corresponding reference dataset (Fig. 3 **c**), validating cfDecon’s ability to capture both condition-specific and cell-type-specific variations in cfDNA signature deconvolution.

We benchmark cfDecon’s cell-type-specific signature prediction performance across diverse conditions. In WGBS analysis, cfDecon achieves an average CCC of 0.953 across six scenarios (Supplementary Table. S6). When applied to the UXM reference dataset, cfDecon reveals distinct beta value patterns for five major blood cell types across five methylation channels (Fig. 3 **a**). Compared to the Baseline-S, cfDecon demonstrates superior performance with 73.2% and 65.6% reductions in L1 scores under fixed and unfixed settings, respectively, while maintaining higher CCC values of 0.7 and 0.59 (Fig. 3 **d**,**e**). The visualization results (Fig. 3 **f**) validate cfDecon’s robust signature prediction capability in both predominant blood cells (e.g. Blood-Granul) and rare cell types (e.g. Endothelium). While CelFEER shows competitive performance, it exhibits overfitting tendencies. Detailed results are available in Supplementary Fig. S3, where it is evident that CelFEER is demonstrating signs of overfitting across all cases. Meanwhile, our cfDecon far outperforms the Baseline-S in terms of performance.

### 3.3 Deconvolution on Clinical Datasets Enables Accurate Disease Diagnosis

To validate cfDecon’s biological relevance, we assess its prediction accuracy against established clinical knowledge using two datasets with documented clinical records and previously reported cell-type composition patterns, ALS and HCC [7,10]. For each disease, we follow a systematic analytical pipeline: first generating comprehensive proportion predictions across all 35 cell types, then focusing specifically on disease-relevant cell types (skeletal muscle cells for ALS and liver hepatocytes for HCC), and finally evaluating the model’s diagnostic potential through ROC curve analysis. This validation strategy demonstrates cfDecon’s ability to capture disease-specific cell-type patterns with potential diagnostic implications.

In evaluating cfDecon’s efficacy for ALS diagnosis, we examine cfDNA samples from ALS patients and healthy controls [7]. Consistent with the known composition of cfDNA, blood cells emerge as the predominant contributors across all samples. Detailed analysis in Fig. 4 **a**. demonstrate a significant elevation of skeletal muscle-derived cfDNA in ALS patients relative to healthy controls. Quantitatively, ALS patients exhibit a mean skeletal muscle cfDNA fraction of 0.0032 ± 0.0028, compared to 0.0005 ± 0.0005 in healthy subjects—a six-fold increase suggesting potential utility as a non-invasive biomarker. Statistical validation through both parametric and non-parametric tests confirms the significance of these findings (rank-sum test, 2.424, p = 0.0153). Furthermore, ROC curve analysis for disease detection reveals superior performance of cfDecon (AUC = 0.79) compared to CelFEER (AUC = 0.53), demonstrating cfDecon’s enhanced capability to detect disease-specific alterations in cfDNA composition that could be valuable for non-invasive diagnostics and disease monitoring (Fig. 4 **b**. and Supplementary Fig. S7). More results of cfDNA fraction predictions across methods for ALS are shown in the Supplementary Table. S5.

**Fig. 4:**
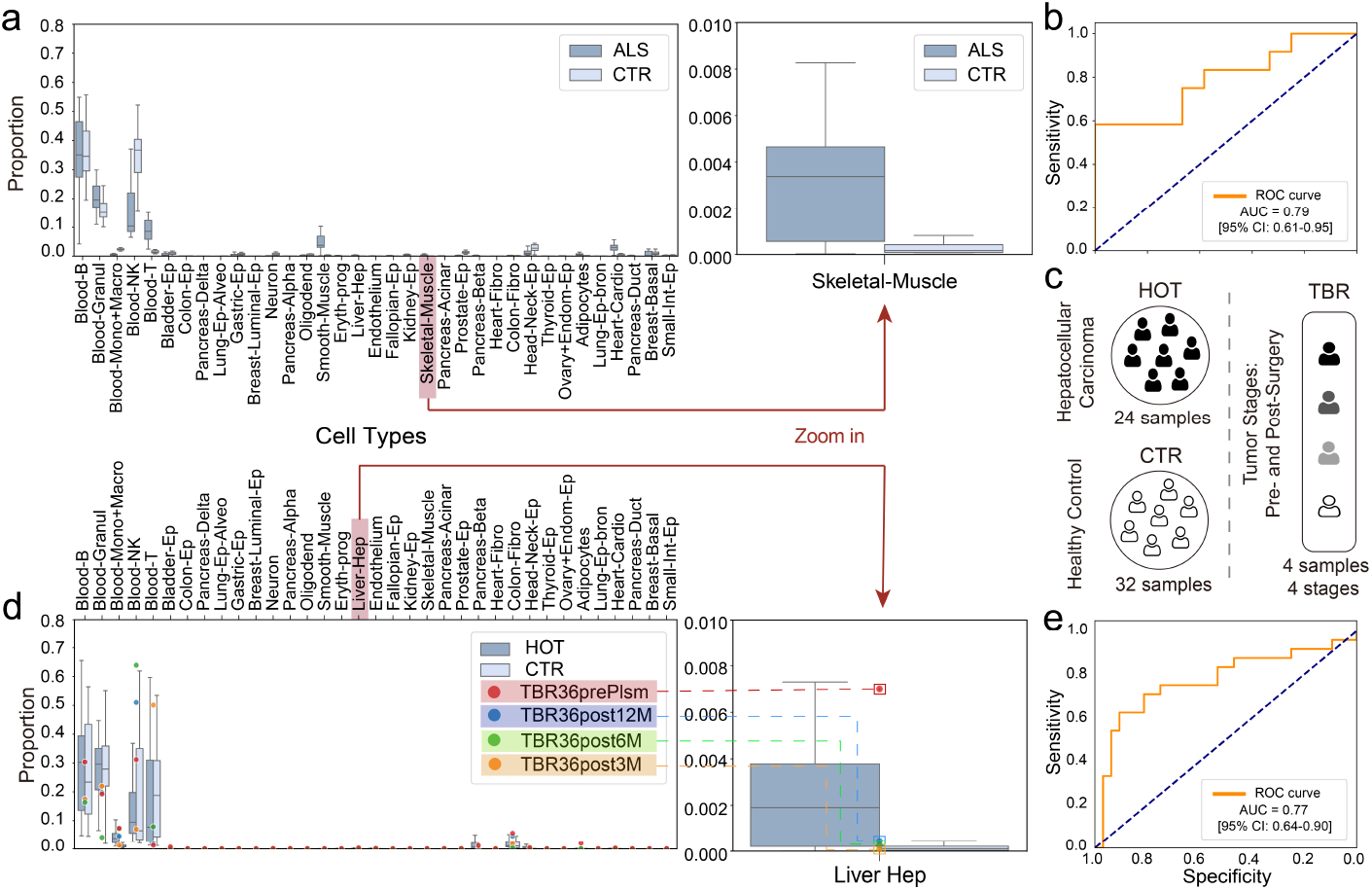
Deconvolution on datasets with clinical information. **a**. Predicted proportions in ALS patients and controls (CTR) with zoom-in on skeletal muscle cells (right). **b**. Receiver operating characteristic (ROC) curves and area under ROC curve (AUC) for ALS detection using the predicted proportion of the disease-relevant cell type. **c**. The illustration of the HCC dataset includes the sample groups of HCC patients (HOT), CTR, and tumor stages (TBR). **d**. Predicted proportions for 35 cell types in HOT and CTR with a zoom-in on liver hepatocyte cell (right), where TBR36 samples represent different time points post-surgery. **e**. ROC curves and AUC for HCC detection using the predicted proportion of the disease-relevant cell type.

Next, we examine cfDecon’s performance in HCC, analyzing cfDNA samples from 24 HCC patients (HOT), 32 healthy controls (CTR), and 4 TBR samples from 2 patients with documented tumor stages (illustration of this dataset can be found in Fig. 4 **c**) [10]. Consistent with expected cfDNA profiles, blood cells constitute approximately 80% of the total composition in both groups. Notably, we observe a significant elevation in liver-derived cfDNA in HCC patients, with mean hepatocyte fractions of 0.0010 ± 0.0011 compared to 0.0002 ± 0.0008 in healthy controls—a five-fold increase in Fig. 4 **d**. Statistical analysis confirm the significance of these findings (rank-sum test, 3.444, p = 0.000574). Furthermore, cfDecon effectively track the post-surgery recovery pattern in TBR36 patients, where liver hepatocyte proportions show an initial decrease from pre-surgery levels (TBR36prePlsm), followed by stabilization at levels comparable to healthy controls through subsequent time points (3M, 6M, and 12M post-surgery), suggesting successful treatment response. ROC curve analysis in Fig. 4 **e**. and Supplementary Fig. S7. demonstrat superior diagnostic performance of cfDecon (AUC = 0.78) compared to CelFEER (AUC = 0.55). More results of cfDNA fraction predictions across methods for HCC are shown in the Supplementary Table. S6. Collectively, these results demonstrate cfDecon’s capability to not only distinguish HCC patients from healthy controls but also track disease progression through treatment, highlighting its potential as a non-invasive monitoring tool.

### 3.4 Sensitive Detection of Disease-related Differentially Methylated CpG Regions

To further demonstrate the clinical utility of cfDecon, we employ a systematic approach to identify the most relevant CpG regions by comparing methylation signatures between disease groups (HCC/ALS) and their corresponding control groups within cell-type-specific contexts (liver hepatocytes for HCC and skeletal muscle for ALS). Following the workflow illustrated in Fig. 5 **a**, we select the top 10-15% of differentially methylated CpG regions and conduct a channel-specific analysis. The analysis results (in Fig. 5 **b**.) demonstrate the strongest influence from high methylation channels, followed by low methylation channels, which aligns with recent epigenetic research showing that epigenetic organization predominantly consists of methylated, unmethylated, and bimodal regions [39], and supports the concept that cell-type-specific methylation typically manifests as unmethylated states against a default methylated background [49]. These selected CpG regions are then visualized in Fig. 5 **c**, where we can observe that cfDecon exhibits higher sensitivity in detecting methylation differences across channels for skeletal muscle cell types compared to CelFEER. The visualization of the global CpG region is shown in Supplementary Fig. S8.

**Fig. 5:**
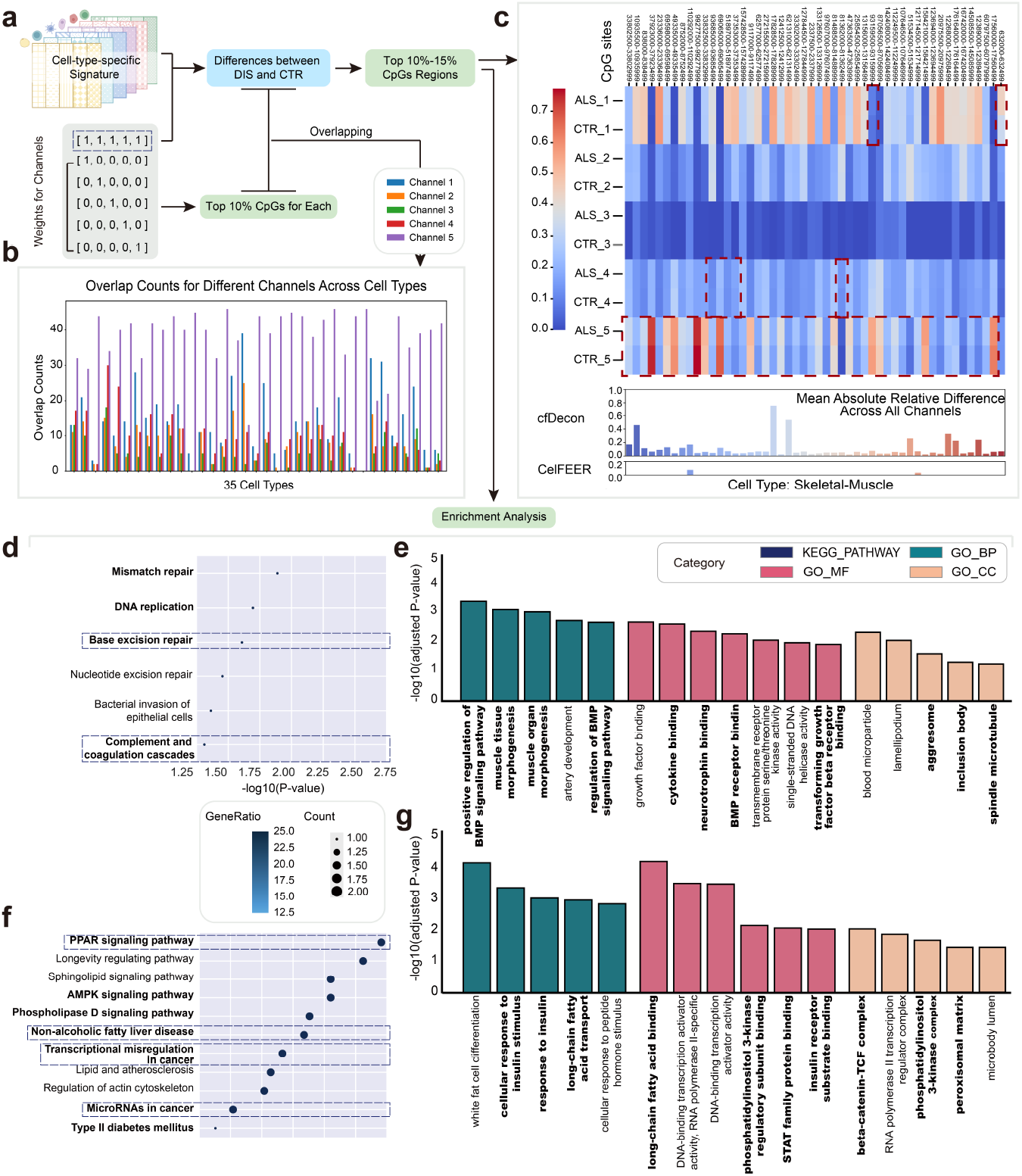
Enrichment analysis of differential CpG region in ALS and HCC. **a**. Workflow for cell-type-specific signatures analysis. Top CpG regions are selected by calculating the differences in cell-type-specific signatures between disease and control groups. The top 10%-15% of differentially methylated CpGs are then further analyzed. **b**. The bar plot displays the overlap counts of the top 10% CpGs across different methylation channel weights for individual cell types, demonstrating the varying influence of each channel on CpG selection. **c**. Heatmap visualization of the selected CpGs in skeletal muscle cells for ALS and control samples across multiple channels, highlighting methylation pattern differences. Below are the bar plots of the mean absolute relative difference across all five channels. **d**,**e**. KEGG pathway and GO enrichment analysis results for ALS, with significantly enriched pathways displayed. **f**,**g**. KEGG pathway and GO enrichment analysis results for HCC, with significantly enriched pathways displayed. The GO enrichment analysis is categorized into Biological Process (BP), Molecular Function (MF), and Cellular Component (CC). Bolded terms indicate pathways of particular relevance to the disease mechanism. All enriched terms are shown in panels **d**-**g**. meet the criteria of p-value < 0.05 and q-value < 0.2, ensuring statistical significance and minimizing false discovery rates in the enrichment analysis.

We then analyze whether the differentially methylated CpG regions are biologically significant using KEGG and GO databases for pathway enrichment. The KEGG pathway analysis of differential methylation patterns in ALS samples reveals significant enrichment in DNA repair pathways (Fig. 5 **d**), including base excision repair, mismatch repair, and DNA replication [38,13,35], suggesting a link between DNA damage repair mechanisms and motor neuron degeneration. The enrichment of complement and coagulation cascade pathways further highlights the role of neuroinflammation in ALS progression [84,15,3]. The GO analysis in Fig. 5 **e**. identifies enrichment in biological processes related to muscle tissue morphogenesis [36,73] and BMP signaling pathway regulation [2,62], molecular functions involving growth factor binding and neurotrophin binding [69,74,17,63], and cellular components including aggresomes and inclusion bodies [12,77,57]. These findings underscore the complex interplay between DNA damage, neuroinflammation, and protein aggregation in ALS pathogenesis, providing potential therapeutic targets for future investigation. Shifting our focus to HCC, the KEGG pathway analysis of significantly differential methylated CpGs between HCC and control samples reveals several crucial pathways in HCC pathogenesis (Fig. 5 **f**.), including transcriptional misregulation in cancer, microRNA involvement [6,72,26], and metabolic disorders such as NAFLD and type II diabetes mellitus [66,59]. The GO analysis in Fig. 5 **g**. further identifies significant enrichment in biological processes related to insulin response and fatty acid metabolism [51,76,56], molecular functions involving PI3K regulatory subunit binding and STAT family protein binding [68,30,80,79], and cellular components including the beta-catenin-TCF complex and peroxisomal matrix [46,24,16,54,61]. These findings highlight the intricate interplay between metabolic dysregulation and transcriptional aberrations in HCC pathogenesis, offering potential therapeutic targets and biomarkers for future research. More detailed enrichment analysis is available in Supplementary S2.

In conclusion, cfDecon effectively identifies disease-specific methylation signatures in ALS and HCC, demonstrating superior performance in detecting cell-type-specific epigenetic alterations. This enhanced performance stems from rigorous methodological controls, including careful prevention of data leakage, ensuring that findings reflect genuine biological signals rather than technical artifacts.

## 4 Discussion

cfDNA emerges as a powerful tool for enabling noninvasive diagnostics and monitoring aberrant methylation. Deciphering its cellular origins remains crucial for understanding tissue-specific epigenetic regulation and disease mechanisms. Current computational methods face significant challenges, including the loss of information from using aggregated CpG site data and the inability to generate condition-aware cell-type-specific profiles. Additionally, existing approaches struggle to handle the complexity of cfDNA mixtures containing numerous cell types. cfDecon’s low-rank structure and iterative refinement process offer a comprehensive solution for analyzing these mixtures. The ability to predict cell-type proportions while generating condition-aware cell-type-specific signatures advances the field, improving our understanding of disease-specific methylation patterns and enhancing diagnostic capabilities in personalized medicine.

Future work will focus on two main directions. For model development, incorporating time-series module to cfDecon could enable better monitoring of disease progression and treatment response. We will incorporate prior biological knowledge about cell type-specific methylation markers and regulatory networks to enhance result interpretability and provide deeper insights into disease mechanisms. For comprehensive validation, we will validate our approach on larger and more diverse datasets, including high-quality cancer datasets from Li et al. and Van et al. [44,18], to evaluate model generalization ability across various disease contexts. We will also explore more challenging scenarios such as early-stage cancers for cfDNA deconvolution.

